# High-accuracy Decoding of Complex Visual Scenes from Neuronal Calcium Responses

**DOI:** 10.1101/271296

**Authors:** Randall J. Ellis, Michael Michaelides

**Affiliations:** National Institute on Drug Abuse Intramural Research Program, Baltimore, MD 21224, United States; Icahn School of Medicine at Mount Sinai, New York, NY 10029, United States; Department of Psychiatry, Johns Hopkins Medicine, Baltimore, MD 21287, United States

**Keywords:** calcium imaging, deep learning, neural networks, neuroscience, Allen Brain Observatory

## Abstract

The brain contains billions of neurons defined by diverse cytoarchitectural, anatomical, genetic, and functional properties. Sensory encoding and decoding are popular research areas in the fields of neuroscience, neuroprosthetics and artificial intelligence but the contribution of neuronal diversity to these processes is not well understood. Deciphering this contribution necessitates development of sophisticated neurotechnologies that can monitor brain physiology and behavior via simultaneous assessment of individual genetically-defined neurons during the presentation of discrete sensory cues and behavioral contexts. Neural networks are a powerful technique for formulating hierarchical representations of data using layers of nonlinear transformations. Here we leverage the availability of an unprecedented collection of neuronal activity data, derived from ∼25,000 individual genetically-defined neurons of the parcellated mouse visual cortex during the presentation of 118 unique and complex naturalistic scenes, to demonstrate that neural networks can be used to decode discrete visual scenes from neuronal calcium responses with high (∼96%) accuracy. Our findings highlight the novel use of neural networks for sensory decoding using neuronal calcium imaging data and reveal a neuroanatomical map of visual decoding strength traversing brain regions, cortical layers, neuron types, and time. Our findings also demonstrate the utility of feature selection in assigning contributions of neuronal diversity to visual decoding accuracy and the low requirement of network architecture complexity for high accuracy decoding in this experimental context.

## Introduction

Understanding how the brain detects, organizes, and interprets information from the external world is a major question in neuroscience and a critical barrier to the development of high-performance brain-computer interfaces and artificial intelligence (AI) systems. At a fundamental level, such efforts rely on understanding how sensory information is encoded by the brain and conversely, how this information can be decoded from brain activity. Among our senses, vision is arguably the most important contributor for interacting with our environment. A variety of technologies have been used to observe and deduce visual encoding from neuronal activity responses (Klimesch, Fellinger, and Freunberger, 2011; Machielsen et al., 2000; Vinck, Batista-Brito, Knoblich, and Cardin, 2015). However, none of these approaches allow the examination of visual encoding in discrete, genetically-defined neurons. Calcium’s presence and role in the nervous system has been studied for decades (Graziani, Escriva, & Katzman, 1965) and was first imaged *in vivo* relatively recently using fluorescent dyes (Stosiek, Garaschuk, Holthoff, & Konnerth, 2003). Genetically-encoded calcium indicators (e.g. GCaMPs) (Nakai, Ohkura, and Imoto, 2001) were later used for monitoring calcium changes in genetically-defined neurons using optical imaging and/or photon detection technologies (Göbel and Helmchen, 2007; Tian et al., 2009). It is now generally accepted that GCaMP activity serves as a valid proxy for real-time imaging of in vivo neuronal activity (Huber et al., 2012; Ohki et al., 2005; Resendez and Stuber, 2015).

Visual decoding from calcium imaging data has a relatively sparse history, with greater prior focus placed on visual decoding of electrophysiological data (Warland, Reinagel, and Meister, 1997; Pillow et al., 2008; for a review, see Quiroga and Panzeri, 2009). Machine learning algorithms, specifically hierarchical neural networks, have been recently developed, that along with matching human performance on object categorization, predicted neuronal responses to naturalistic images in two areas of the ventral stream in nonhuman primates (Yamins et al., 2014). Other studies have also reported impressive performance using conventional (e.g., linear) machine learning architectures. In one study, intracranial field potentials in patients with intractable epilepsy were recorded while images were presented (Quiroga, Reddy, Koch, and Fried, 2007), and mean decoding accuracy across 32 images was reported at 35.4%, with chance being 3.1% (1/32). In another similar study (Liu, Agam, Madsen, and Kreiman, 2009), binary classification accuracy of ∼95% was achieved, with ∼60% classification accuracy using five classes. In nonhuman primates, single-trial classification accuracies of 82-87% were reported (Manyakov, Vogels, Van Hulle, 2010), and in another study primary visual cortical responses to 72 classes of static gratings were decoded with 60% accuracy (Graf, Kohn, Jazayeri, and Movshon 2011). More recently, perfect decoding accuracy using dorsomedial ventral stream data from nonhuman primates was achieved in a five-class image recognition task (Filippini et al., 2017). Importantly, while some of these prior studies reported high, and in one case, perfect decoding accuracy, none of these prior studies achieved high accuracy with a high number of classes.

In vivo neuronal calcium imaging, while requiring substantial video and other downstream processing (Harris, Quiroga, Freeman and Smith, 2016; Peron, Chen, and Svoboda, 2015), enables delineation of neuronal traces using fluorescent signals from discrete, genetically-defined neurons over time, without having to employ simulations. To date, calcium activity has been used to visually decode movie scenes with high accuracy from small numbers of high-responding neurons using nearest mean classification (Kampa, Roth, Göbel, and Helmchen, 2011). In particular, Kampa et al. selected high-responding neurons based on correlations between responses in a single trial to other trials for both individual neurons and neuronal populations. In machine learning terminology, this is a biologically-inspired form of feature selection, where specific features are chosen to make a model more parsimonious, easier to interpret, and less likely to overfit. Simple linear classification of calcium responses from larger (∼500) populations of neurons was also recently implemented using natural and phase-scrambled movies (Froudarakis et al., 2014). This work demonstrated that total activation of primary visual cortical neurons does not differ between anesthesia and wakefulness, but that population sparseness is heightened during the latter. Froudarakis et al. also showed that this phenomenon enables more accurate visual decoding. Importantly, these prior studies used small numbers of visual stimuli and employed small numbers of recorded neurons. Notably, while the former of the two studies achieved high decoding accuracy, the probability of accurate decoding by random chance was high (i.e., 25%). To our knowledge, high visual decoding accuracy using many unique and complex visual stimuli has not been previously reported.

The implementation of deep neural networks has proven successful for high-accuracy visual classification of images using features such as skin lesions (Esteva et al., 2017), facial recognition (Li et al., 2015), and for deducing the brain’s physiological age from MRI scans (Cole et al., 2016). However, deep neural networks have not been applied yet to visual decoding using calcium responses. In most visual classification tasks using deep learning, inputs are images, where classifiers are trained on examples of different image classes and then used to classify a validation set of images from these same classes. In the context of deep learning using calcium imaging data, the inputs are not images but neuronal calcium responses *to* images. Accordingly, for such data, classifiers are trained on responses *to* sensory stimuli and consequently are labeled *by* the sensory stimulus. In this way, when incorporating diverse sources of responses (e.g., brain regions, neuron populations), the differential classification accuracy between these sources can serve as an indicator for how well they process information individually but also collectively as an integrated circuit. Here, the unique advantage of calcium imaging over other modalities is the unique ability to make observations and answer physiological questions by distinguishing specific neuronal types at the individual and population levels. As such, imaging neuronal calcium responses in behaving animals enables the investigation of discrete neurons, neuronal populations, whole brain regions, and brain circuits while retaining the neuron as the fundamental unit composing all these echelons.

Exploiting advances in instrumentation, software, genetic engineering, and viral vector-based genetic targeting technologies, the Allen Institute for Brain Science recently published an extensive data set of neuron-specific GCaMP6 activity measures (http://observatory.brain-map.org/visualcoding; Hawrylycz, et al. 2016). In particular, 597 experiments were conducted using mice from transgenic Cre recombinase-expressing lines co-expressing GCaMP6 in six genetically-defined neuronal types across six regions of the visual cortex (primary (VISp), anterolateral (VISal), anteromedial (VISam), lateral (VISl), posteromedial (VISpm), and rostrolateral (VISrl)) and eleven cortical depths (175, 265, 275, 300, 320, 325, 335, 350, 365, 375, 435 µm). GCaMP6 activity, as a function of neuron type, region, and depth, was measured in response to the presentation of several types of visual stimuli. Stimuli included natural scenes, static gratings, drifting gratings, and movie clips. The data collection and analysis methods used for the Allen Brain Observatory (ABO) dataset are available in a whitepaper (Allen Institute for Brain Science, 2017). Raw calcium data along with various corrections for tens of thousands of neurons are made available through the Allen Institute’s software development kit (SDK).

Here, we describe the application of supervised machine learning to decoding visual scenes using data from the ABO. We first trained four different machine learning architectures on calcium responses to presentation of 118 unique natural scenes, and then tested the ability of these models to classify calcium responses based on the presented scene. All training and validation was performed in a frame-by-frame manner, beginning with the scene preceding the scene of interest (proximal scene), and training on every frame through the two subsequent (distal) scenes. That is, models were trained and validated on all frames composing the scene preceding the proximal scene (prior scene), the proximal scene itself, the first distal scene, and the second distal scene. Each model was trained on calcium responses from neuronal populations distinguished by neuron type, brain region, and cortical depth in response to all 118 scenes. Further, calcium responses were also distinguished by their response properties in two different ways. The first was a biologically-informed feature selection technique where neurons were selected if they showed a positive mean response across all visual scenes. The second was a conventional feature selection technique, employing different numbers of neurons with the highest ANOVA F-values for the target labels.

## Methods

### Data collection and segmentation

Using the ABO SDK (accessed in June 2016), calcium traces were segmented by region, neuron type, and cortical depth. Protocols showing the time points of visual scene presentation were retrieved for each experiment to label corrected (Δ*F/F*) GCaMP6 traces recorded from ∼25,000 neurons of the visual cortex in response to 118 unique natural scenes. In each experiment, calcium responses were measured from individual genetically-defined neurons, in a subregion of the visual cortex, at a single cortical depth. Session-long calcium traces from all individual neurons (**Fig. 1A**) were segmented by 118 natural scenes each shown 50 times in random order, for a total of 5900 scene presentations. For each experimental condition (i.e., all experiments corresponding to a combination of neuron, region, and cortical depth), a three-dimensional array (Walt, Colbert, and Varoquaux, 2011) was generated where rows, columns, and ranks corresponded to scenes, neurons, and frames respectively (**Fig. 1B**). Because each of the 118 natural scenes was presented 50 times in each experiment, all arrays had 5900 rows. The number of neurons (i.e. columns) varied by experimental condition, but all arrays captured 28 frames, making 28 ranks in the third dimension. These 28 frames represented the 7 frames of the prior scene, the 7 frames of the scene used to label the trace (proximal scene), and the two subsequent, “distal” scenes (**Fig. 1C**). 40 presentations of each scene were used for training and 10 for validation, yielding an 80/20 split where 4720 total presentations were used for training and 1180 for validation. Separate models were trained and validated for each of the 28 frames. For example, at frame 1, networks were trained on 4720 calcium traces at frame 1 and then validated on 1180 calcium traces at frame 1. Each of these traces represented all the neurons in the respective experimental condition at the selected frame. This process was repeated for all 28 frames.

**Figure 1.**
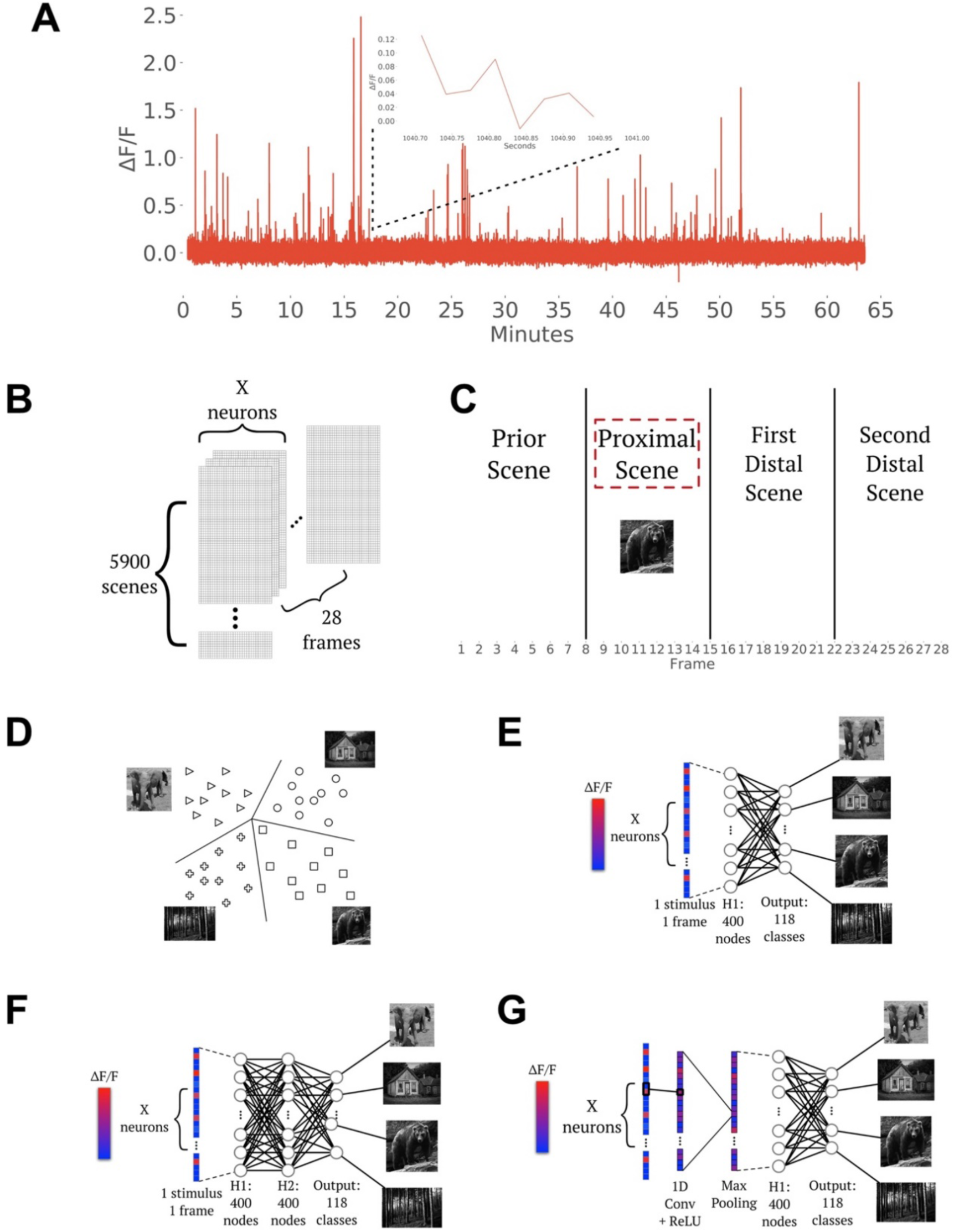
Data organization and network architectures. (**A**) Single representative neuron GCaMP6 trace over a ∼63-minute session. (**B**) 3D arrays were constructed where rows, columns, and ranks corresponded to scenes, neurons, and frames respectively. (**C**) Temporal breakdown of scenes and frames. Architectures utilized: (**D**) support vector machine (SVM), (**E**) single hidden-layer neural network (SNN), (**F**) two hidden-layer neural network (DNN), and (**G**) convolutional neural network (CNN).

### Architectures

We tested four machine learning architectures on calcium trace classification that were implemented in Scikit-learn (Pedregosa et al., 2011) and Keras (Chollet, 2015): a support vector machine (SVM), a shallow neural network (SNN), a deep neural network (DNN), and a convolutional neural network (CNN). Training was conducted for 20 epochs at each frame and evaluated at the corresponding frame in the validation set. All networks used an 80/20 train/validation split, with 4720 visual responses at a single frame for training and 1180 responses for validation.

A SVM (**Fig. 1D**) was implemented in Scikit-learn using the OneVsRestClassifier function with a linear support vector classifier and a regularization parameter calculated using grid search. A SNN (**Fig. 1E**) consisted of one batch normalization layer (Ioffe and Szegedy, 2015), a dropout layer (0.5) (Srivastava et al., 2014), a flattening layer, one dense layer with a rectified linear (relu) activation function and 400 nodes, a dropout layer (0.5), and a final dense layer with 118 nodes (for 118 classes) with a softmax activation function. An Adam optimizer was used for adjustment of learning rates (Kingma and Ba, 2014). The DNN (**Fig. 1F**) consisted of one batch normalization layer, one hidden fully connected layer, dropout, another fully connected layer, another dropout, and a fully connected output layer. Finally, a CNN (**Fig. 1G**) was tested which consisted of one batch normalization layer, a 1D convolution, 1D MaxPooling, flattening, a dense layer (relu), dropout, and a final dense output layer. Categorical cross-entropy was used to measure loss in trace classification and accuracy was used to quantify correct classifications.

### Neural Population Comparisons

#### Visual Cortex Decoding Accuracy as a Function of Neuron Type and Cortical Depth

We measured decoding accuracy for all neurons in each of the six visual cortical regions, ignoring differences of neuron type and cortical depth. In addition, to control for effects due to the number of neurons within a given region, decoding accuracy was further measured for each of the six regions after limiting the number of neurons by the lowest number imaged within a single region (VISam, 1514 neurons). For the five regions containing more than 1514 neurons, 1514 neurons were randomly selected. This enabled a comparison of all regions in terms of visual decoding accuracy without the confound of differing numbers of imaged neurons.

For the highest resolution of neuronal population segmentation, we measured decoding accuracy for all neuron types in all regions at all cortical depths, for a total of 63 populations. We then compared populations by randomly limiting each to 250 neurons, retaining 32 datasets. This number was selected to retain the greatest number of datasets for population comparison while using a population size in range of previously published calcium imaging experiments (Barnstedt et al., 2015; Lecoq et al., 2014).

#### Visual Cortex Decoding Accuracy as a Function of Biologically-inspired and Conventional Feature Selection

Visual decoding accuracy was measured in each population using only neurons with a high average response (Δ*F/F* > 0.01) across all 5900 scene presentations at any of the latter 21 frames, during the proximal scene and the two distal scenes. In each population, only neurons with a mean Δ*F/F* response higher than 0.01 across all scene presentations within a single frame, from the beginning of the proximal scene to the end of the second distal scene, were used for training and validation. We refer to these neurons as *high mean responders* (HMRs).

We used a univariate feature selection technique implemented in Scikit-learn (SelectKBest with the default ANOVA F-test) to select the top 10, 50, 100, 250, 500, 750, 1000, 1250, and 1500 neurons in each region at each frame for visual decoding. Our feature selection was run only on the training data, and the validation data from the corresponding neurons were used accordingly.

## Results

We first assessed decoding accuracy for each of the four machine learning architectures on a frame-by-frame basis in six regions of the mouse visual cortex using all imaged neurons in each region across 28 frames. The highest decoding accuracy achieved was in VISp (94.66%) using calcium responses from 8661 neurons as input to a CNN (**Figs. 2A-C, Table S1**). While the SNN, DNN and CNN all achieved similar peak accuracies, the SVM performed noticeably worse across all regions. Looking at changes in accuracy across frames for all regions, accuracy began to increase at frames 10-11 (the proximal scene began at frame 8), continued increasing after the proximal scene ended and the first distal scene began (frame 15), reached peak accuracy at frame 18 and stayed above chance throughout the duration of the two distal scenes (frames 14-28).

**Figure 2.**
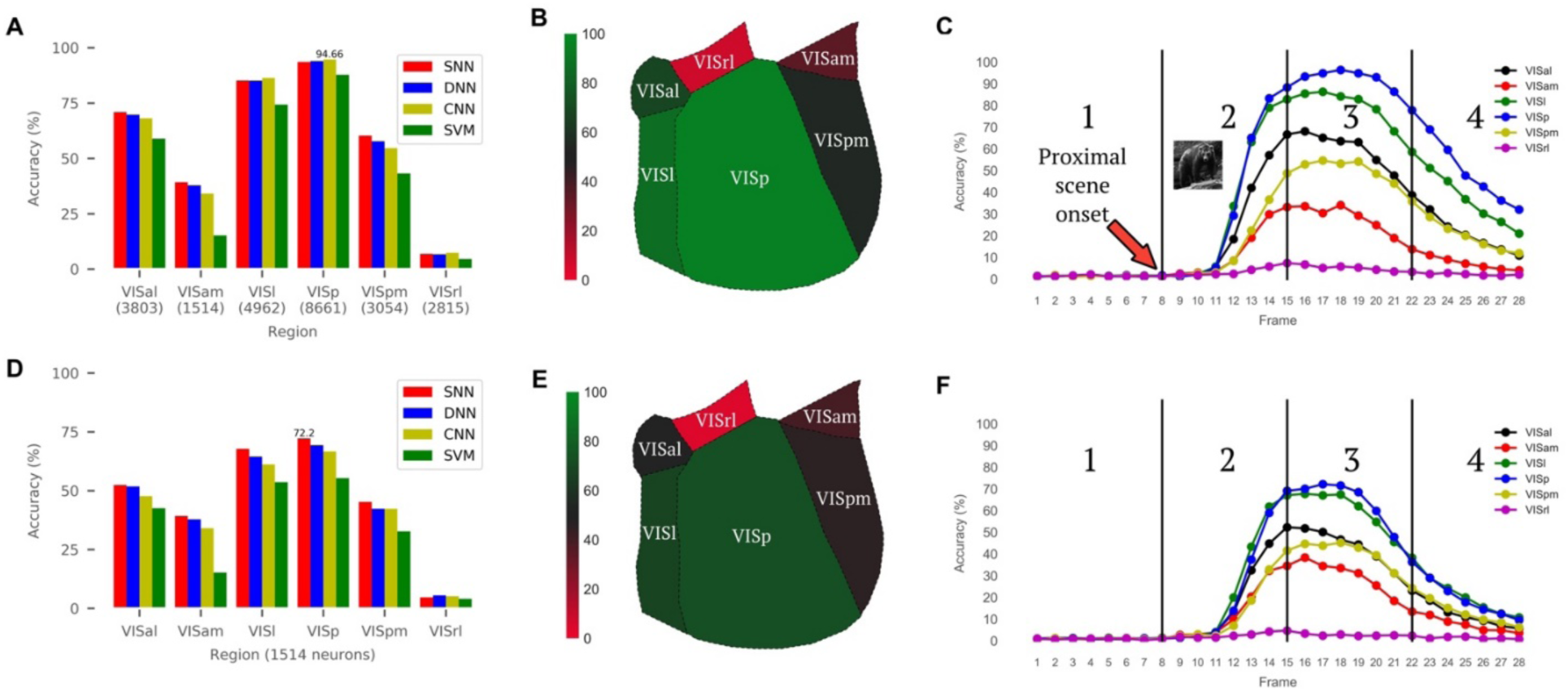
Decoding accuracy for six regions of the mouse visual cortex. (**A**) Peak accuracies across all frames for four different machine learning architectures. (**B**) Heatmap plot overlaid onto a horizontal view of the mouse visual cortex indicating cortical subregions as a function of accuracy (0-100%) using a CNN; data from (**A**). (**C**) Frame-by-frame accuracies for each region when decoding was performed using a CNN. Scene 1 refers to scene presented prior to the scene that the trace is labeled by. Scene 2 is the proximal scene, (the scene the trace is labeled by and the one being decoded). Scenes 3 and 4 are the two distal scenes presented after the proximal scene. (**D**) Peak accuracies across all frames for four machine learning architectures are shown when neuronal inputs for each region were limited to 1514 randomly-chosen neurons. (**E**) Heatmap plot overlaid onto a horizontal view of the mouse visual cortex indicating cortical subregions as a function of accuracy (0-100%) using a SNN; data from (**D**). (**F**) Frame-by-frame accuracies for each region when decoding was performed using a SNN. Scene classification as described in (**C**).

To assess the differences in visual decoding accuracy between the six regions of the mouse visual cortex and to control for the number of neuronal inputs, we limited the number of neurons analyzed in each region by the lowest number of total neurons in any of the six regions (VISam, 1514 neurons). We included all 1514 neurons from VISam and then randomly chose 1514 from each of the other five regions and calculated decoding accuracies for all regions across all 28 frames. As above, VISp showed the highest accuracy (72.2%) compared to all other regions (**Figs. 2D-F, Table S2**), albeit, at notably lower levels than previously when a larger number of inputs were used (**Figs. 2A-C, Table S1**). In contrast to the prior comparison, the highest accuracy was observed at frame 17 (one frame earlier) and using the SNN. Overall, for both approaches, the regions were ranked as follows in descending order of accuracy: VISp, VISl, VISal, VISpm, VISam, and VISrl and all regions, with the exception of VISrl exhibited a similar frame-by-frame pattern of accuracy.

Next, we assessed visual decoding accuracy as a function of cell type, cortical depth and region, for a total of 63 populations (**Fig. 3A, Table S3**). Using all available neurons in each population, the highest decoding accuracy achieved was 77.97% with Cux2-expressing neurons in VISp at a depth of 175µm using a SNN (**Fig. 3B**). This specific neuron type exhibited the highest accuracy among all other populations and a distinct frame-by-frame accuracy profile which was shifted to the right compared to the rest of the populations examined (**Fig. 3C**). Cux2-expressing neurons at 175 µm depth in VISp showed peak accuracy at frame 18 whereas other Cux2-expressing neurons at 275 µm depth in either VISp or VISl showed peak accuracy at earlier frames (frames 15, 16), a difference of about 30 ms. Rorb- and Emx1-expressing neurons showed peak accuracy at frames 15 and 16 respectively (**Fig. 3C**). Notably, three out of five of the top performing populations were Cux2-expressing neurons. Additionally, four out of the top five performing populations originated in VISp. For all five populations, the SNN performed best of the four tested architectures.

**Figure 3.**
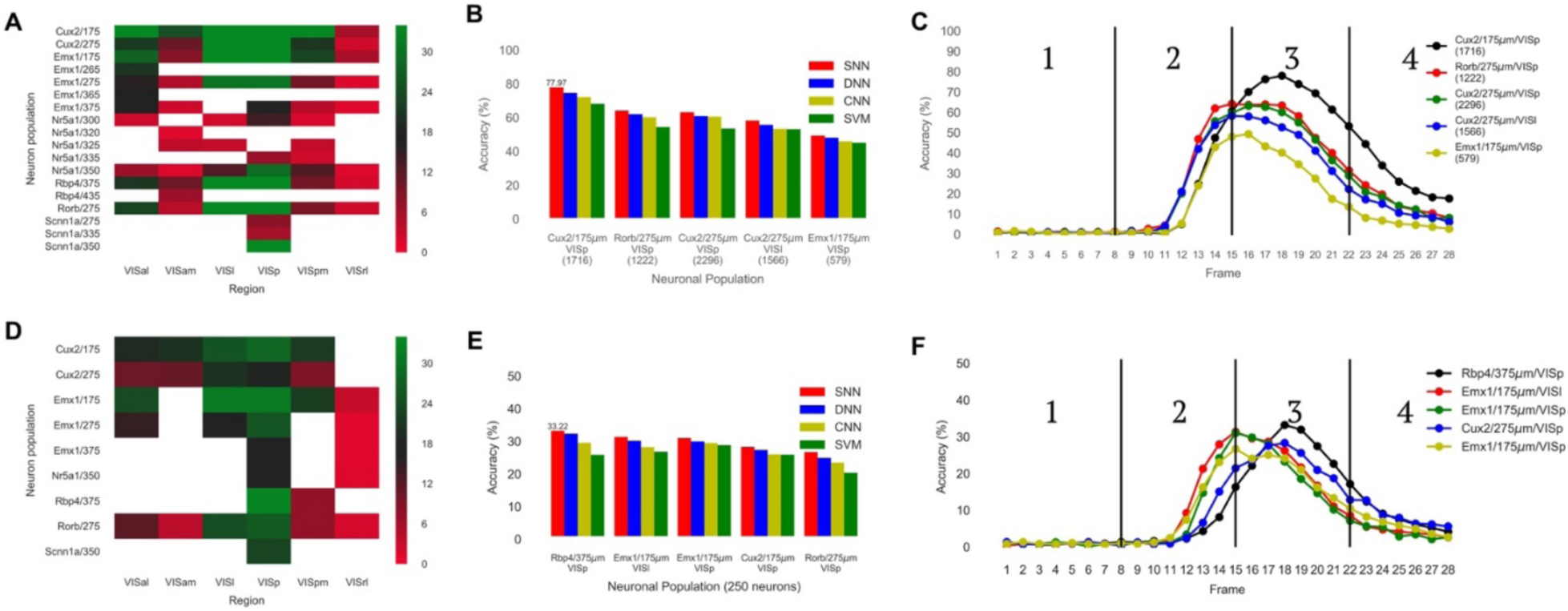
Decoding accuracy for the top neuronal populations parcellated by region, neuron type, and cortical depth. (**A**) Peak accuracies across all frames for four machine learning architectures are shown. **(B)** Peak accuracies across all frames for four machine learning architectures are shown when limited to 250 randomly-chosen neurons.

Next, we limited each of these populations to 250 randomly-selected neurons (32 populations, with the other 31 containing less than 250 total neurons) (**Fig. 3D, Table S4**). In this dataset, Rbp4 neurons in VISp at 375µm showed the highest decoding accuracy of 33.22% at frame 18 using the SNN (**Figs. 3E, F**). Again, four out of the five best performing populations were derived from VISp. As above, neuron types differed in the time-course for peak accuracy. Emx1-expressing neurons showed peak accuracy at frame 15, which did not depend on depth or cortical subregion (**Fig. 3F**). In contrast, Rbp4- and Cux2-expressing neurons within VISp but at different depths, exhibited peak accuracy at frame 18 (**Fig. 3F**).

In all six regions of the visual cortex, we measured visual decoding accuracy as a function of neuronal response using HMRs: neurons that showed a mean response greater than a value of 0.01 Δ*F/F* across all 5900 scene presentations in any of the latter 21 frames (proximal scene and two distal scenes). We found that accuracy was greater than or within 3% of the accuracy when using all neurons in the respective region (**Figs. 4A, B, Table S5**). The highest accuracy achieved was at frame 18 in VISp using the CNN (**Fig. 4C**). To explore the differences in accuracy between HMRs and other neurons (non-HMRs (nHMRs)), we compared identically-sized samples of HMRs and nHMRs (583 of each) with a SNN in all regions. This number was chosen based on the minimum number of total HMRs contained in any of the six regions (VISam). We found that the accuracy of HMRs was 1.5-3-fold greater than that of nHMRs (**Figs. 4D, E, Table S6**) with the highest accuracy observed in VISp at frame 18 (**Fig. 4F**). Next, we assessed decoding accuracy in HMRs as a function of region, neuron type, and depth. All samples of HMRs showed similar or higher peak accuracies compared to randomly-selected neurons by between 3-6%. To compare HMRs and nHMRs in these parcellated populations, we were forced to make comparisons of 35 neurons each due to many populations having small numbers of either HMRs or nHMRs. Nevertheless, even with very sparse populations of neurons, the accuracies of HMRs maintained their 1.5-3-fold greater level than those of nHMRs (**Figs. 4G, H, Table S7**). As above, the highest frame accuracy was observed in Rbp4-expressing neurons at 375 µM in VISp and at frame 19 (**Fig. 4I**).

**Figure 4.**
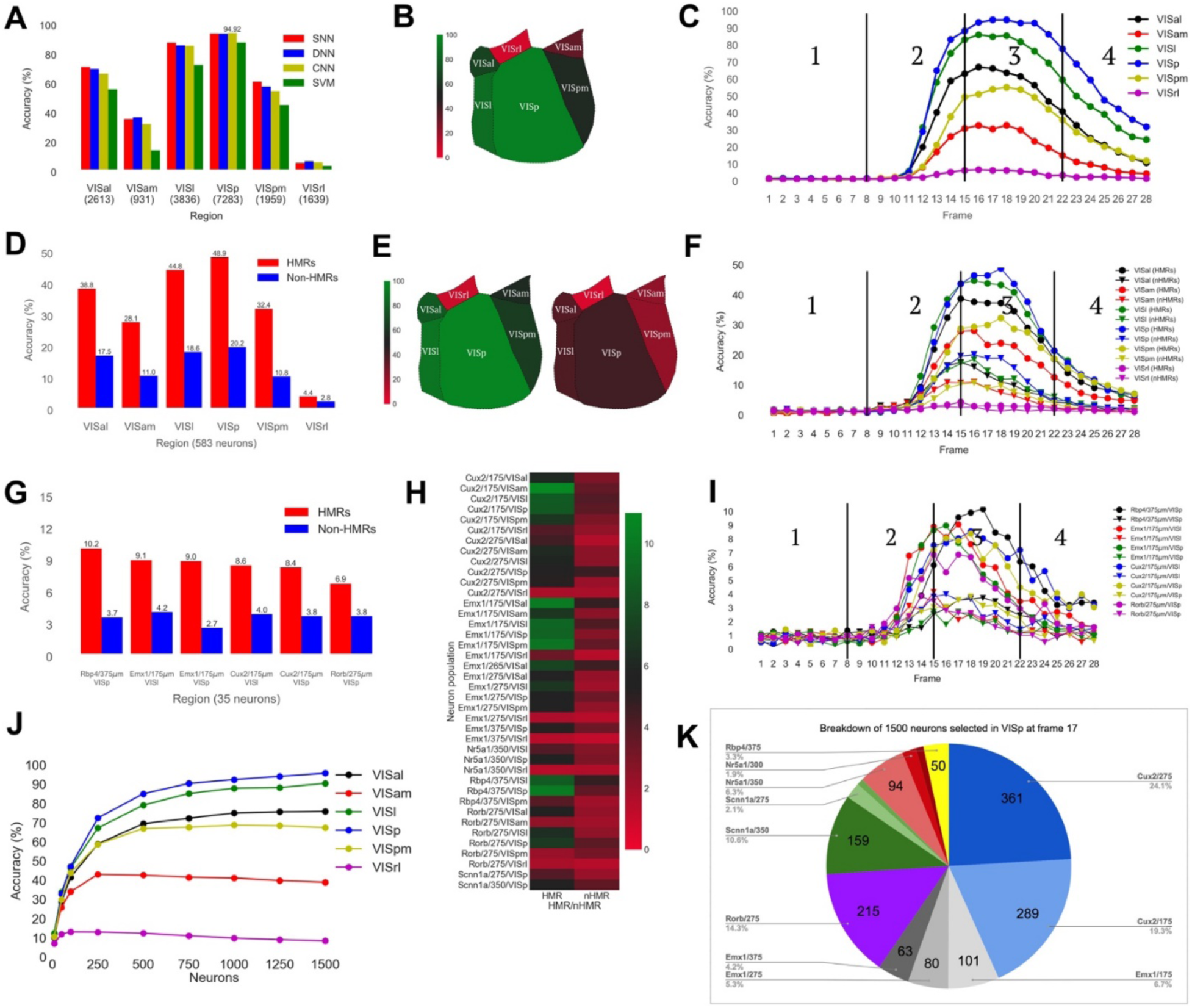
Decoding accuracy for the top neuronal populations parcellated by region, neuron type, and cortical depth selected after biologically-inspired and feature classifications. **(A)** Peak accuracies across all frames for four machine learning architectures are shown when neurons for each region were limited to high mean responding neurons. **(B)** Peak accuracies across all frames for a SNN are shown when limited to 583 high mean responding and 583 non-high mean responding neurons. **(C)** Peak accuracies across all frames for a shallow neural network are shown when limited to 35 high mean responding and 35 non-high mean responding neurons. **(C)** Peak accuracies across all frames for a shallow neural network are shown when neurons for each region were limited to feature selected neurons (**G**) Breakdown of the 1500 feature selected neurons in VISp during the frame where peak accuracy was achieved.

Finally, in each recorded region of visual cortex, an F-test was conducted, and the *k* best neurons were selected for visual decoding. Values of *k* were 10, 50, 100, 250, 500, 750, 1000, 1250, and 1500. Using the 1500 neurons with the highest F-values in VISp, an accuracy of 95.76% was achieved, the highest of all experiments (**Fig. 4J, Table S8**). Critically, groups of neurons selected by F-value performed better than either the totality of the neurons in each region, or all the HMRs in each region. In all cases, feature-selected populations were much smaller than the total HMRs and total neurons in these regions. Across all visual scenes and neuron types, the SNN performed about as well, and in some cases better than the CNN. For natural scene prediction in VISp, all networks reached accuracies of 85-95%. However, the highest accuracy achieved across all experiments (95.76%) was obtained using 1500 neurons selected by F-value in VISp. VISp showed the highest accuracies for neurons selected by F-test, mean response, and when limiting all regions to the same number of randomly-selected neurons. A specific breakdown of neuron classification at the highest accuracy achieved using 1500 neurons in shown in **Fig. 4K**.

## Discussion

Here we describe the application of supervised machine learning to visual decoding of neuronal calcium responses to 118 unique and complex naturalistic scenes. Our findings describe a neuroanatomical map of decoding accuracy in the mouse visual cortex in response to complex naturalistic scenes and as a function of regional cortical parcellation, depth, and neuron type. A general finding was that, regardless of neuron type or cortical depth, the highest visual decoding accuracy was achieved in VISp while the lowest was achieved in VISrl. This observation is consistent with prior findings from a recent study that used this same dataset (Esfahany, Siergiej, Zhao, and Park, 2017) and known information regarding the hierarchical organization of the mouse visual cortex (Glickfeld, Reid, and Andermann, 2014). Of note, for almost all populations, accuracy remained above chance (1/118, 0.85%) throughout the duration of the two scenes distal to the scene that the calcium response was labeled by. Additionally, we found that both feature selection methods we utilized enabled similar, or in some cases, higher accuracies compared to retaining all neurons in the respective population, indicating that different forms of feature selection can reduce the processing load of decoding algorithms without compromising, and even sometimes improving, decoding accuracy. Of note, the accuracy of some regions declined when adding more neurons by highest F-value (i.e., VISam, VISpm, and VISrl). This was most apparent for VISrl, where peak accuracy was achieved with only 100 neurons, and all other groups including more neurons performed worse, indicating that there may exist a threshold where increasing the number of such neurons introduces noise to the data, making the accuracy dwindle.

A novel key finding of our study was the capability of a SNN architecture to achieve high-accuracy visual decoding using this type of data, especially in the context of the many classes included. Taken together with prior work that used a CNN trained on ImageNet (Deng et al., 2009) to decode both seen and imagined visual stimuli from fMRI data (Horikawa & Kamitani, 2017), our findings indicate that visual discrimination can be modeled effectively using data spanning different imaging modalities and across species. It is important to point out, however, that the highest decoding accuracy achieved in that prior study was ∼80% for binary classification, an accuracy level considerably lower than what we report here for the number of classes included. Since fMRI is expected to be less proximal to neuronal activity than *in vivo* calcium imaging, our findings support the notion that high-resolution imaging modalities which capture information closer to base neuronal physiology may be more effective at reaching higher levels of decoding accuracy, especially when using simple architectures.

Another notable finding was that the highest decoding accuracy (95.76%) was achieved by utilizing a neuronal population selected using a conventional feature selection approach, the F-test. This was specifically observed within VISp at frame 17 during the presentation of the first distal scene to the scene decoded. Interestingly, this high accuracy level was achieved using a population containing a diverse set of neuron types at various depths within this region (**Fig. 4K**). Indeed, looking at all the neurons in VISp during the 21 frames beginning with the proximal stimulus and extending through the second distal stimulus, for a total of 31500 neurons (1500×21), the top three most represented populations were Cux2/275µm (7881 neurons), Cux2/175µm (6007), and Scnn1a/350µm (2744). When ignoring depth, the most represented population was Cux2 (7881 + 6007 = 13888; 13888/31500 = 44.09%), followed by Emx1 (4952 neurons, 15.72%), and Rorb (4476 neurons, 14.21%). This contrasts with the breakdown of VISrl, the worst-performing region, where the top three populations, accounting for depth, were Nr5a1/350µm (9116), Emx1/275µm (5480), and Emx1/175µm (3514). When not accounting for cortical depth, the top three performing populations were Emx1 (12165 neurons), Nr5a1 (9116), and Cux2 (4622). Overall, Cux2 was the most represented neuron type in the best-performing region and composed 44% of feature-selected VISp neurons but only 15% of feature selected VISrl neurons. Cux2 is reported to be a critical regulator of dendritic branching, spine development and synapse formation in cortical layer 2/3 neurons (Cubelos et al., 2010). However, looking at populations segmented by region, depth, and neuron type, Rbp4/375µm/VISp was overall the best-performing population. The Allen Institute has profiled these and other genetically-defined neurons to build a comprehensive taxonomy of the adult mouse cortex (Tasic et al., 2016). Referring to this data, five out of six of the above studied neuron types are excitatory (with no information available for Emx1), and thus likely to project to other cortical areas or subcortical regions. Importantly, while we did observe differences in decoding accuracy between the six different genetically-defined neuron types sampled in VISp, the top three performing classes of neurons never differed more than 5% from each other. Interestingly, this difference was observed using calcium responses from randomly-selected neurons as well as randomly-selected HMRs. Collectively, these findings indicate that neuronal diversity within the visual system hierarchy plays a key role in decoding accuracy, but ultimately it is the visual system regional hierarchy that is the main contributor. Regarding the contribution of cortical depth to the accuracy signal, while we did observe differences between different neuronal populations, these were relatively small, further supporting the notion that most of the variation in visual decoding accuracy was accounted for by neuron location within visual cortical regions, as opposed to neuron type and depth.

Another important finding was that in all experiments where models performed above chance, decoding accuracy peaked ∼210-360 ms after the presentation of a given scene, a time-point that coincided with the presentation of the first distal scene. Interestingly, above-chance decoding accuracy was maintained over the duration of two distal scenes across many of the neuronal populations investigated. Why the accuracy consistently peaked during the presentation of a distal scene is unclear, though we hypothesize it may represent a delay in calcium dynamics or the optimal imaging methods used to record them. In previous work on visual decoding of categories from human magnetoencephalography data, decoding accuracy peaked 80-240ms after stimulus onset (Carlson, Tovar, Alink, and Kriegeskorte, 2013) and decayed over the period of one second. Each image was shown for 533 ms, meaning the accuracy peaked during the presentation of the proximal image. Additionally, after stimulus presentation in this prior study, a delay period between 900-1200 ms in length was given. This means the duration of the decay in accuracy occurred within the window of the stimulus presentation and subsequent delay period. In contrast, in the current study, accuracy peaked 210-360 ms after scene onset, or 0-150 ms after the onset of the first distal scene and continued for an additional 240-390 ms. We propose the term *refractory processing* to denote this delayed temporal property of calcium in allowing the decoding of visual scenes during the subsequent presentation of a unique stimulus. While we cannot say exactly what this phenomenon represents, the appearance of this property in visually-evoked calcium dynamics may be related to the recently discovered phenomenon of perceptual echo in human occipital EEG responses to changes in luminance (Chang et al., 2017). Importantly, our finding also agrees with the findings of Filippini et al. where neural activity during the delay after object presentation yielded greater decoding accuracy than the object presentation itself (2017). Like Carlson et al.’s study, the difference between our findings and those of Filippini et al. is rather than a delay after object presentation, different scenes were continuously presented after one another, meaning time points with the highest decoding accuracy coincided with the presentation of another scene, rather than a lack of stimulus.

Finally, we found that regional decoding accuracy was maintained or improved beyond using all of a given region’s neurons by limiting the selection of neurons to those with mean responses above 0.01 Δ*F/F* to all presentations of naturalistic scenes (independent of the type/content of the scene) at any frame between the onset of the proximal stimulus through the duration of the two distal stimuli. For all neuronal populations, when including only HMRs for decoding, accuracy either exceeded or was within 3% of the accuracy compared to when all neurons within that same population were included. Further supporting this was the stark difference in decoding accuracy between number-matched samples of HMRs and non-HMRs, where HMRs performed 1.5-3x better than nHMRs. Additionally, using a conventional feature selection technique from machine learning, accuracy was maintained or improved using even fewer neurons than those selected by mean response. These observations indicate that visual decoding accuracy is strongly determined by the response properties of discrete neurons within the visual system hierarchy. From a biological perspective, this suggests that complex and diverse visual imagery, independent of content, may be collectively encoded in this discrete population of neurons with high response profiles to visual image presentation, or neurons that simply show strongly differentiated responses to visual images. For clarity, we assert that neuronal diversity plays an important role in visual decoding, but what seems to be even more important is the regional hierarchy of the visual system that produces such distinct performances in decoding accuracy.

In sum, here we describe a neuroanatomical map of the mouse visual cortex decoding aptitude of different regions and neuron types at various cortical depths and shed light on the temporal dynamics of visual encoding and decoding using a neural network approach as they persist across the presentation of a large and diverse collection of complex visual scenes. Our findings demonstrate the low requirement of neural network architecture complexity in the context of visual decoding using neuronal calcium data and highlight the strong contributions of regional localization, neuronal response profile, the quantity of recorded neurons, discrete genetically-defined neuronal populations and cortical depths to visual information encoding and visual decoding. Additionally, the temporal trajectory of decoding accuracy throughout the duration of scene presentations indicated that accuracy peaked roughly 300 ms after the scene appeared, during the presentation of a unique stimulus, an observation we refer to as *refractory processing*, that may reflect an inherent property of neurons in the visual cortex. Finally, we show that feature selection techniques from machine learning can parse out neuronal populations most indicative of differentiated responses to complex naturalistic scenes and increase decoding accuracy.

A limitation of this study is the small numbers of neurons in some of the parcellated populations. Parcellating the data by region, neuron type, and depth sometimes yielded populations with less than five neurons, which had little value in assessing decoding accuracy. If these populations had contained hundreds or thousands of neurons, we could have perhaps seen with greater clarity how those specific parcellated populations of neurons would compare to the others and within themselves in terms of a fixed number of randomly selected neurons, HMRs and nHMRs. This leaves open questions about the functional importance of these parcellated populations in comparison to others within the same region. We plan to revisit this in the future as more data from the ABO becomes available. In future work we also plan to better understand the features of the calcium signal which our networks accurately differentiated in the context of many classes and small number of examples and furthermore, how calcium responses from other brain systems perform in this context.

## Acknowledgments

This work was funded by the National Institute on Drug Abuse Intramural Research Program (ZIA000069). MM is a cofounder and owns stock in Metis Laboratories.

---

**Supplementary Table 1.** Peak accuracies for all regions and machine learning architectures, the frames these accuracies were achieved in, and the number of neurons used for decoding in each region.

| **Regional decoding accuracies, all neurons** |  |  |  |  |
| --- | --- | --- | --- | --- |
| **Brain Region** | **SNN** | **DNN** | **CNN** | **SVM** |
| **VISal**<br>**3803 neurons** | 70.93%<br>15 | 69.75%<br>16 | 68.14%<br>16 | 58.9%<br>17 |
| **VISam**<br>**1514 neurons** | 39.24%<br>16 | 37.8%<br>15 | 34.07%<br>16 | 15.17%<br>16 |
| **VISl**<br>**4962 neurons** | 85.17%<br>16 | 85.08%<br>16 | 86.36%<br>17 | 74.32%<br>17 |
| **VISp**<br>**8661 neurons** | 93.56%<br>17 | 93.98%<br>17 | **94.66%**<br>**18** | 87.8%<br>18 |
| **VISpm**<br>**3054 neurons** | 60.25%<br>18 | 57.71%<br>16 | 54.66%<br>17 | 43.22%<br>18 |
| **VISrl**<br>**2815 neurons** | 6.86%<br>15 | 6.69%<br>15 | 7.37%<br>15 | 4.49%<br>16 |

**Supplementary Table 2.** Peak accuracies for all regions limited to 1514 neurons for each machine learning architecture, and the frames these accuracies were achieved in.

| **Regional decoding accuracies, limited to 1514 neurons** |  |  |  |  |
| --- | --- | --- | --- | --- |
| **Brain Region** | **SNN** | **DNN** | **CNN** | **SVM** |
| **VISal** | 52.37%<br>15 | 51.77%<br>16 | 47.71%<br>16 | 42.54%<br>16 |
| **VISam** | 39.24%<br>16 | 37.8%<br>15 | 34.07%<br>16 | 15.17%<br>16 |
| **VISl** | 67.8%<br>16 | 64.49%<br>17 | 61.19%<br>17 | 53.64%<br>18 |
| **VISp** | **72.2%**<br>**17** | 69.32%<br>18 | 66.69%<br>17 | 55.34%<br>17 |
| **VISpm** | 45.25%<br>18 | 42.29%<br>18 | 42.29%<br>18 | 32.71%<br>18 |
| **VISrl** | 4.66%<br>15 | 5.51%<br>15 | 5.17%<br>15 | 4.07%<br>15 |

**Supplementary Table 3.** Peak accuracies for the top five neuronal populations parcellated by region, neuron type, and cortical depth, for each machine learning architecture, and the frames these accuracies were achieved in.

| **Neuron type/region/depth decoding accuracies, all neurons** |  |  |  |  |
| --- | --- | --- | --- | --- |
| **Population (Top 5)** | **SNN** | **DNN** | **CNN** | **SVM** |
| **Cux2, VISp, 175µm,**<br>**1716 neurons** | **77.97%**<br>**18** | 74.66%<br>18 | 72.12%<br>18 | 68.22%<br>19 |
| **Rorb, VISp, 275µm,**<br>**1222 neurons** | 64.15%<br>15 | 62.03%<br>16 | 60.25%<br>16 | 54.41%<br>17 |
| **Cux2, VISp, 275µm,**<br>**2296 neurons** | 63.22%<br>16 | 60.93%<br>16 | 60.59%<br>16 | 53.47%<br>16 |
| **Cux2, VISl, 275µm,**<br>**1566 neurons** | 58.22%<br>15 | 55.76%<br>15 | 53.39%<br>16 | 53.14%<br>16 |
| **Emx1, VISp,**<br>**175µm,**<br>**579 neurons** | 49.32%<br>16 | 48.14%<br>15 | 45.93%<br>16 | 45.08%<br>16 |

**Supplementary Table 4.** Peak accuracies for the top five neuronal populations parcellated by region, cell type, cortical depth, and limited to 250 neurons. Accuracies are shown for each machine learning architecture, and the frames they were achieved in.

| **Neuron type/region/depth decoding accuracies, limited to 250 neurons** |  |  |  |  |
| --- | --- | --- | --- | --- |
| **Population (Top 5)** | **SNN** | **DNN** | **CNN** | **SVM** |
| **Rbp4, VISp,**<br>**375µm** | **33.22%**<br>**18** | 32.37%<br>18 | 29.58%<br>18 | 25.85%<br>18 |
| **Emx1, VISl, 175µm** | 31.36%<br>15 | 30.17%<br>15 | 28.22%<br>15 | 26.78%<br>15 |
| **Emx1, VISp,**<br>**175µm** | 31.02%<br>15 | 30%<br>15 | 29.49%<br>15 | 28.81%<br>15 |
| **Cux2, VISp,**<br>**175µm** | 28.31%<br>17 | 27.37%<br>18 | 26.02%<br>17 | 25.93%<br>17 |
| **Rorb, VISp, 275µm** | 26.69%<br>15 | 24.92%<br>15 | 23.47%<br>17 | 20.34%<br>17 |

**Table 5.** Peak accuracies for all regions limited to high mean responding neurons for each machine learning architecture, and the frames these accuracies were achieved in.

| **Regional decoding accuracies, all high mean responders** |  |  |  |  |
| --- | --- | --- | --- | --- |
| **Brain Region** | **SNN** | **DNN** | **CNN** | **SVM** |
| **VISal**<br>**2613 neurons** | 71.78%<br>16 | 70.34%<br>16 | 67.12%<br>16 | 56.44%<br>18 |
| **VISam**<br>**931 neurons** | 36.27%<br>16 | 37.46%<br>15 | 32.88%<br>18 | 14.75%<br>18 |
| **VISl**<br>**3836 neurons** | 88.14%<br>17 | 86.53%<br>16 | 86.19%<br>16 | 73.05%<br>17 |
| **VISp**<br>**7283 neurons** | 94.49%<br>17 | 94.41%<br>18 | **94.92%**<br>**18** | 88.31%<br>18 |
| **VISpm**<br>**1959 neurons** | 61.78%<br>18 | 58.39%<br>18 | 55.25%<br>18 | 45.76%<br>19 |
| **VISrl**<br>**1639 neurons** | 6.36%<br>15 | 7.46%<br>15 | 6.78%<br>16 | 4.32%<br>17 |

**Table 6.** Peak accuracies for all regions limited to 583 high mean responding and non-high mean responding neurons for a shallow neural network, and the frames these accuracies were achieved in.

| **Regional decoding accuracies, 583 HMRS**<br>**vs. 583 nHMRS, shallow neural network** |  |  |
| --- | --- | --- |
| **Brain**<br>**Region** | **583 HMRS** | **583 nHMRS** |
| **VISal** | 38.81%<br>15 | 17.54%<br>15 |
| **VISam** | 28.13%<br>16 | 11.02%<br>16 |
| **VISl** | 44.83%<br>16 | 18.56%<br>16 |
| **VISp** | **48.9%**<br>**18** | 20.17%<br>16 |
| **VISpm** | 32.37%<br>18 | 10.76%<br>16 |
| **VISrl** | 4.41%<br>15 | 2.80%<br>13 |

**Table 7:** Peak accuracies achieved with feature selected neurons across six regions of the mouse visual cortex and comparison to total HMRs and total neurons in each region. Each cell lists the peak accuracy, the number of neurons in the group, and the frame the accuracy was achieved in.

| **Neuron type/region/depth decoding accuracies, 35 HMRs vs. 35 nHMRs, SNN** |  |  |
| --- | --- | --- |
| **Population (Top 6)** | **35 HMRs** | **35 nHMRs** |
| **Rbp4, VISp, 375µm** | **10.17%**<br>**19** | **3.73%**<br>**19** |
| **Emx1, VISl, 175µm** | **9.07%**<br>**17** | **4.15%**<br>**15** |
| **Emx1, VISp, 175µm** | **8.98%**<br>**16** | **2.71%**<br>**15** |
| **Cux2, VISl, 175µm** | **8.56%**<br>**18** | **3.98%**<br>**16** |
| **Cux2, VISp, 175µm** | **8.39%**<br>**18** | **3.81%**<br>**18** |
| **Rorb, VISp, 275µm** | **6.86%**<br>**15** | **3.81%**<br>**14** |

**Table 8.** Peak accuracies for the top six performing neuronal populations limited to 35 high mean responding and non-high mean responding neurons for a shallow neural network, and the frames these accuracies were achieved in.

| **Regional decoding accuracies, Neurons selected by F-score, mean response, and total neurons** |  |  |  |
| --- | --- | --- | --- |
| **Brain Region** | **F-test (SNN)** | **HMRs** | **Total neurons** |
| **VISal** | 75.93%, 1500, 15 | 71.78%, 2613, 16, SNN | 70.93%, 3803, 15, SNN |
| **VISam** | 43.14%, 250, 15 | 37.46%, 931, 15, DNN | 39.24%, 1514, 16, SNN |
| **VISl** | 90.59%, 1500, 15 | 88.14%, 3836, 17, SNN | 86.36%, 4962, 17, CNN |
| **VISp** | **95.76%, 1500, 17** | 94.92%, 7283, 18, CNN | 94.66%, 8661, 18, CNN |
| **VISpm** | 68.81%, 1000, 17 | 61.78%, 1959, 18, SNN | 60.25%, 3054, 18, SNN |
| **VISrl** | 13.22%, 100, 14 | 7.46%, 1639, 15, DNN | 7.37%, 2815, 15, CNN |

